# Beadex modulates hematopoiesis and innate immune defense against microbial infection in *Drosophila melanogaster*

**DOI:** 10.1101/2025.10.13.681973

**Authors:** Sakshi Jain, Kashish Salian, Vrushti Shah, Upendra Nongthomba

## Abstract

Transcription factors play a crucial role during hematopoiesis with implications in leukemia. LIM-domain only proteins (LMO) are also found to be associated with chromosomal translocation in T-cell acute lymphoblastic leukemia and are highly expressed in acute myeloid leukemia, where they correlate with poor prognosis. However, their role in myeloid cell lineage specification and function is poorly understood. Using *Drosophila melanogaster*, which possesses a single LMO homolog, *Beadex* (dLMO), and a conserved myeloid-like immune system, we have shown its conserved role in plasmatocytes (equivalent to macrophages) development and innate immunity. *Beadex* is expressed in progenitor and differentiated plasmatocytes, and its loss reduced lymph gland size, mature plasmatocyte numbers and phagocytic efficiency. The phagocytes of *Beadex* mutant flies feature reduced filopodia length and lamellipodium area. Mechanistically, *Beadex* regulates actin remodelling by regulating the expression level of *profilin* (*chickadee*/*chic),* as *chic* overexpression rescued the defect in phagocytosis. Overexpression of *Beadex* in hemocytes increased fly susceptibility towards *Salmonella enterica serovar* Typhimurium infection and this mortality was due to an infection-induced pathology independent of bacterial burden. Bulk RNA sequencing of *Beadex* knockdown hemocytes revealed transcriptional changes in actin cytoskeleton regulatory genes, phagosome function and immune response pathways. In conclusion, our study identifies *Beadex* as a key transcriptional regulator impacting multiple aspects of myeloid cell function, ranging from hematopoiesis and cytoskeletal dynamics to host-level immune defense.

## Introduction

Transcription factors form highly regulated and tightly organised networks regulating tissue development both spatially and temporally (Zerella et al., 2023). Hematopoiesis is a complex process wherein the hematopoietic stem cells differentiate into the hematopoietic progenitor cells, which subsequently give rise to mature cells of myeloid and lymphoid lineages(Chapman & Zhang, 2023). Network of various transcription factors regulates these cell fate decisions including cell differentiation and lineage specification. Identifying and studying these key players and networks help us to understand how normal hematopoiesis works and how genetic aberrations prevent their functions, leading to haematological malignancies (Zerella et al., 2023).

One such important transcription factor identified in genome wide binding assays as a key regulator of the hematopoietic stem and progenitor cells (HSPC) is LIM-only protein 2 (LMO2), a member of LIM-domain family of proteins(Zerella et al., 2023). This network also includes other transcriptional factors such as FLI1, ERG, GATA2, RUNX1, LYL1 and SCL/TAL1, which regulate HSPC function and are common mutational targets in leukemia, wherein progression to the malignancy is regulated by additional secondary events targeting multiple hematopoietic pathways(Zerella et al., 2023). Aberrant expression of LMO2 is strongly linked to haematological cancers. In T-cell acute lymphoblastic leukemia (T-ALL), LMO2 overexpression resulting from chromosomal translocations, retroviral insertions, or deletions near the LMO2 locus, disrupts T-cell differentiation (Latchmansingh et al., 2022). Loss-of-function studies have indicated that LMO2 is essential for maintaining T-cell progenitors and for the progression of T-cell differentiation program(Hirano et al., 2021). It has been found to be a part of the transcriptional complex involving several DNA binding and adapter proteins like TAL1/SC1, E2A, GATA1, and Ldb1(Hirano et al., 2021). In diffuse large B-cell lymphoma (DLBCL), LMO2 plays an important role in DNA repair as its overexpression drives genome instability by impairing DNA repair, and in acute myeloid leukemia (AML), it is found to be highly expressed and is correlated with reduced survival rates in these patients wherein, in complex with LIM-binding proteins (LDB1) it regulates cell proliferation and prevent apoptosis (Latchmansingh et al., 2022; Lu et al., 2023b). However, the precise role of LMO2 in regulating myeloid cell specification and immune function remains poorly understood.

*Drosophila melanogaster* provides a genetically tractable model to study immune cell function, as it possesses a simplified yet conserved myeloid-like immune system(Zerella et al., 2023;, Evans et al., 2003;, Yu et al., 2022). Three major cell types perform cellular immune functions in Drosophila: plasmatocytes, crystal cells, and lamellocytes, which resemble vertebrate myeloid lineage cells. Plasmatocytes, which make up 90–95% of circulating hemocytes, act as macrophage-like cells, phagocytosing microbes and apoptotic debris while also contributing to antimicrobial peptide (AMP) secretion(Melcarne et al., 2019;, Yu et al., 2022). Crystal cells, comprising about 5% of hemocytes, mediate melanization for wound healing and pathogen defense(Evans et al., 2003a), while lamellocytes are induced during parasitic infections. Importantly, *Drosophila melanogaster* expresses a single LMO homolog, *Beadex* (dLMO), providing a simplified system to investigate LMO protein function in vivo.

*Beadex* is one of the classic Drosophila mutations discovered by Bridges in 1923 and described by Morgan, Bridges, and Sturtevant in 1925. *Beadex* is a dominant mutation that maps to region 17C on the X chromosome (FlyBase Gene Report: Dmel\Bx). The mutant phenotype was found to be beaded wings (long, narrow, and excised edges). *Beadex* was later identified as LIM-only protein (dLMO), a member of the LIM-domain protein family, that plays roles in transcriptional regulation (Milán et al., 1998a; Smith et al., 2014; Zeng et al., 1998). Beyond its well-characterized role in wing development, *Beadex* has also been implicated in immune regulation. It is named a “pathogenesis-specific gene” as its mRNA levels are reduced when infected with a virulent strain of *Pseudomonas aeruginosa* (Apidianakis et al., 2005)*. Beadex* also regulates the immunodeficient (Imd) signalling pathway by controlling the expression of the *Dredd* caspase, thereby influencing AMP production. Additionally, *Beadex* modulates cellular immunity in *Drosophila*, wherein misexpression of *Beadex* led to an altered number of crystal cells, a type of blood cells in Drosophila. The Pnr-Ush complex was discovered as a negative regulator of crystal cell differentiation, and Beadex regulates the levels of the Pnr-Ush complex(Chatterjee et al., 2019). In addition, in genome-wide screens for genes affecting phagocytosis in the *Drosophila* S2 cell line (derived from embryonic plasmatocytes or macrophages), *Beadex* was identified as essential for the phagocytosis process. This study reported that over 90% of cells with *Beadex* knockdown failed to phagocytose *Candida albicans*. Importantly, over 60% failed to engulf non-pathogenic *E. coli* or latex beads(Stroschein-Stevenson et al., 2005). However, Beadex’s in-vivo roles in phagocytosis, and host defense remain unexplored.

Phagocytosis requires rapid remodelling of the actin cytoskeleton to form protruding membrane structures, such as pseudopods and dorsal ruffles, that help in making contact with the target and their subsequent engulfment(Rougerie et al., 2013). The actin remodelling required for forming the phagocytic cup is regulated by actin-binding proteins, such as Arp2/3 complex proteins, myosin motor proteins, and formins(Barger et al., 2022). Despite the identification of *Beadex* in a genetic screen for phagocytosis regulators, the precise mechanism by which it modulates phagocytic efficiency in immune cells remains unclear.

Here, we investigate the role of *Beadex* in Drosophila hematopoiesis and immunity. In this study, using a *Drosophila melanogaster Beadex* hypomorphic and hypermorphic mutants and Gal4/UAS driven RNA interference (RNAi)-mediated knockdown and overexpression in plasmatocytes, we demonstrate that *Beadex* is required to prevent premature differentiation of plasmatocytes, regulates *profilin* (*chic)* expression to support actin remodelling and phagocytosis, and modulate host immune response to bacterial infection. Thus, our findings uncover a previously uncharacterized role for LIM-only protein, *Beadex,* in coordinating transcriptional program with cytoskeleton dynamics in myeloid cell biology.

## Results

### 1. *Beadex* is expressed in both circulating and lymph gland hemocyte populations

Hematopoiesis in Drosophila occurs in two phases (Evans et al., 2003b). The first phase begins during embryogenesis, giving rise to hemocytes, majority of which are plasmatocytes that persist through the larval stages. In the second phase, the lymph gland, a hematopoietic organ, develops during the larval stages, giving rise to adult hemocytes. To assess *Beadex* expression in plasmatocytes derived from different lineages, we used the *Bx-Gal4* driver and a FLAG-HA dual-tagged *Beadex* and a *UAS-GFP* fly line.

In hemocytes expressing *UAS*-*GFP* under *Bx-Gal4*, GFP signal is distributed throughout the cytoplasm confirming uniform expression (Figure 1A & D). Consistent with its predicted role as a transcriptional regulator, FLAG-and HA tagged Beadex showed predominant nuclear localization in circulating hemocytes (Figure 1B & C) and lymph gland cells (Figure 1E). Moreover, Beadex expression is detected in both the progenitor cells in the secondary lobe and in the differentiated as well as progenitor cells of the primary lobe of the lymph gland (Figure 1E). We also confirmed the specificity of FLAG tagged Beadex flyline using *He-Gal4* and *Cg-Gal4* drivers via immunoblotting and observed the band at the predicted size of Beadex, i.e., 42 kDa (Figure 1F).

**Figure 1.**
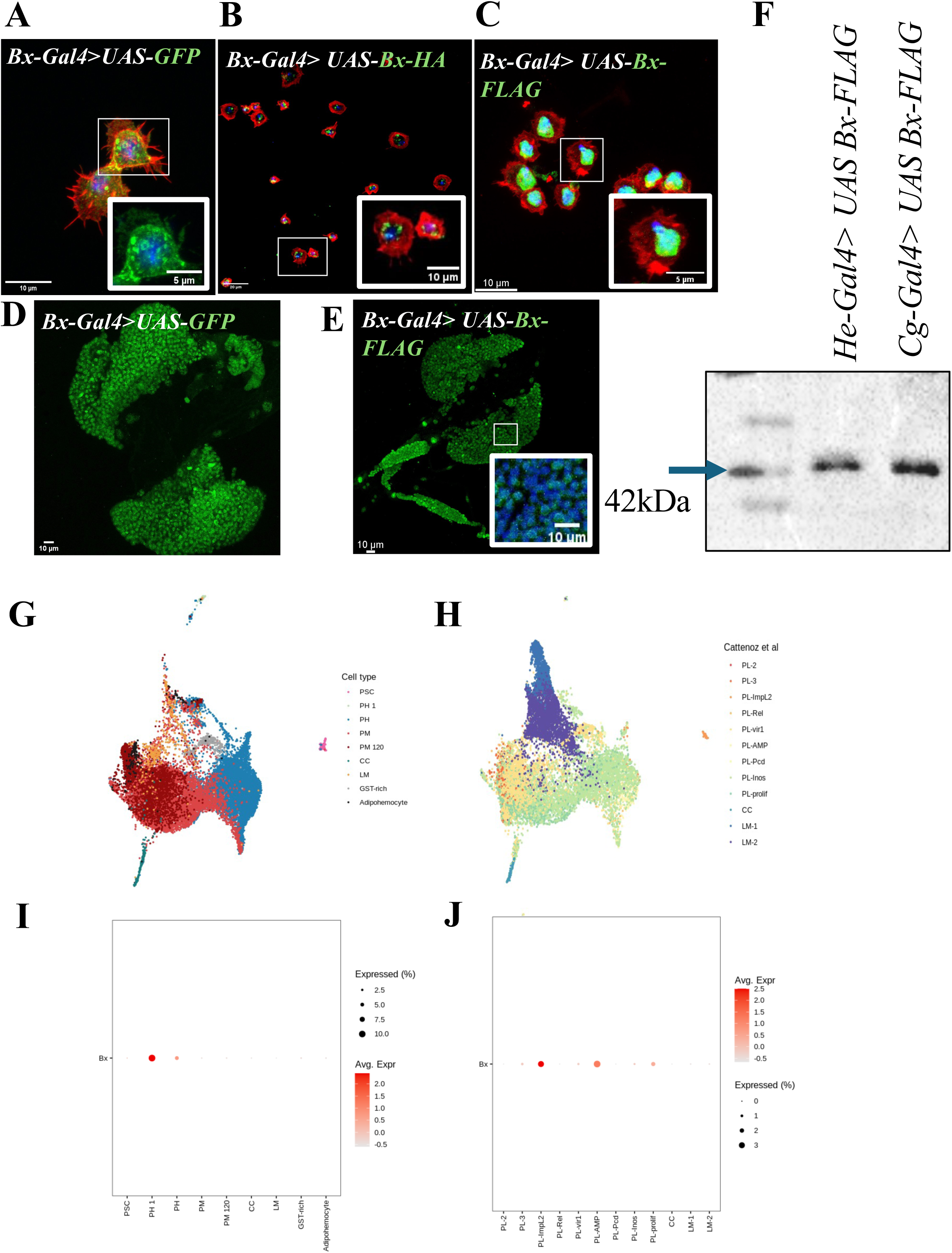
*Beadex* is expressed and localizes to the nucleus in Drosophila hemocytes. Immunofluorescence images showing nuclear localisation of tagged Beadex protein in Drosophila hemocytes (above panel) and lymph gland (lower panel). **A)** Hemocytes are stained with GFP (green) in *Bx-Gal4> UAS-GFP* and with, **B)** HA or **C)** FLAG antibody (green) in overexpressed HA/FLAG tagged Beadex (*Bx-Gal4> UAS-BxRB FLAG-HA*). Cells with counterstained with phalloidin (red) and Hoechst (blue). **D)** Lymph gland are stained with GFP (green) in *Bx-Gal4> UAS-GFP* and with, **E)** FLAG antibody (green) in overexpressed HA/FLAG tagged Beadex (*Bx-Gal4> UAS-BxRB FLAG-HA*). **F)** Western blot analysis of hemocyte lysates from flies overexpressing FLAG-tagged *Beadex* using either the *He-Gal4* or *cg-Gal4* driver. The blot confirms the expression of the Beadex-FLAG protein at the predicted size of ∼42 kDa. **G) & H)** Uniform Manifold Approximation and Projection (UMAP) plot of single-cell RNA sequencing data showing different hemocyte cell clusters. **I) & J)** The dot plot below shows the expression of *Beadex* across the identified cell types, indicating high expression in the prohemocyte cluster and several plasmatocyte populations (http://big.hanyang.ac.kr/flyscrna).

We have also analyzed a published single-cell RNA sequencing database from the FlyscRNAseq portal (http://big.hanyang.ac.kr/flyscrna) (Yoon et al., 2023). Uniform Manifold Approximation and Projection (UMAP) plot of single-cell RNA sequencing data shows different hemocyte cell clusters in lymph gland (Figure 1G) and larval circulating hemocytes (Figure 1H). In the lymph gland, *Beadex* is highly expressed in the cluster of prohemocytes (PH) in the medullary zone (Figure 1I) and in several subpopulations of circulating plasmatocytes in larvae, which include (Figure 1J):

1. PL3 (nearly 9.11% of the total hemocyte population and express genes related to phagocytosis, defense response to bacteria, and extracellular matrix structural constituents).
2. PL-ImpL2 (less than 0.3% of total hemocytes and expresses several transcription factors and markers related to glutathione metabolism).
3. PL-AMP (0.5% of hemocytes that express markers related to the Imd pathway).

Thus, Beadex is expressed in specific subpopulations of circulating larval hemocytes and showed predominant nuclear localization. In addition, it is enriched in the prohemocyte cluster in the lymph gland.

### 2. *Beadex* mutants have a reduced number of plasmatocytes in different developmental stages

To investigate the role of *Beadex* in plasmatocyte development, we analyzed *Bx^7^*, a hopout mutant of *Beadex* generated in our lab, which is a severe hypomorph and exhibits reduced *Beadex* expression in hemocytes (Figure 2A). We observed a significant reduction in plasmatocyte numbers in both larval and adult stages in *Bx^7^* mutants (Figure 2B & C).

**Figure 2.**
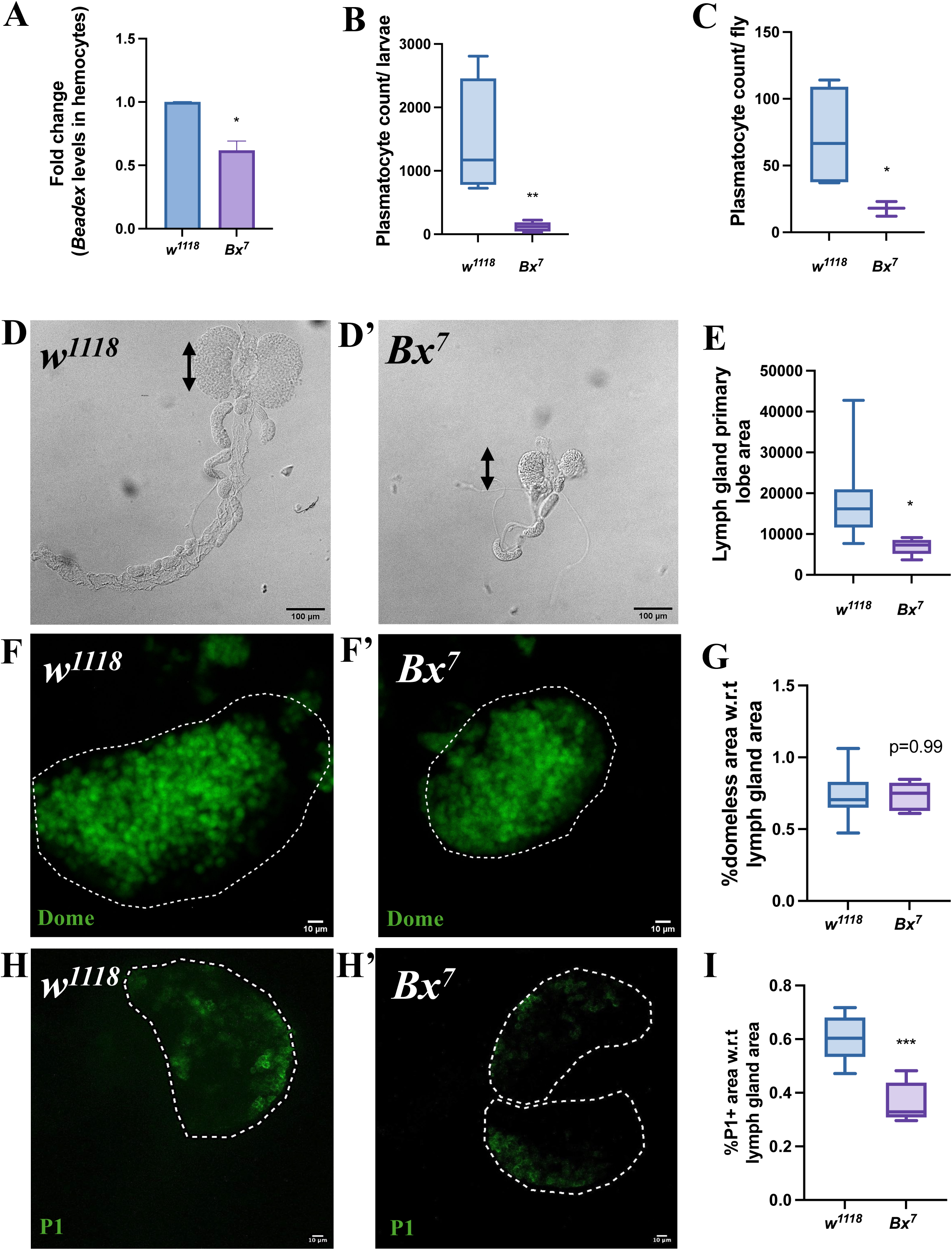
*Beadex* is required for normal plasmatocyte development and differentiation. **A)** Relative transcript levels of *Beadex* in *w^1118^*control and *Bx^7^* mutant larvae, confirming the reduced expression of *Beadex* in hemocytes. Quantification of plasmatocyte number in **B)** larvae and **C)** adult fly, showing a reduction. Brightfield images of larval lymph glands from **D)** *w¹¹¹⁸* and **D’)** *Bx⁷* mutants, showing a significant reduction in the primary lobe (demarcated by white dotted line) area of the mutant. **E)** The bar chart on the right quantifies this reduction. Scale bar, 100 µm. Immunofluorescence images showing the progenitor cell population (marked by Domeless-GFP) in the lymph glands of **F)** *w¹¹¹⁸* and **F’)** *Bx⁷* larvae. **G)** The bar chart on the right quantifies the percentage of the lymph gland area that is Domeless-positive, showing no significant change in the mutant. Scale bar, 10 µm. Immunofluorescence images showing the differentiated plasmatocyte population (marked by P1-positive cells) in the lymph glands of **H)** *w¹¹¹⁸* and **H’)** *Bx⁷* larvae. **I)** The bar chart on the right quantifies the percentage of the lymph gland area that is P1-positive, showing a significant reduction in the mutant. Scale bar, 10 µm. The graph represents the mean of three independent biological replicates and 10 flies are analysed in each case. Statistical significance is denoted by asterisks: *p < 0.05, **p < 0.01, ***p < 0.001.

To access whether the reduction in plasmatocyte numbers reflect a developmental defect, we examined the lymph gland. Our findings indicate that compared to *w^1118^* control the area of the primary lobe is reduced in *Bx^7^*mutants (Figure 2D & E). Expression of Domeless-GFP, a marker of progenitor cells in the medullary zone, showed comparable Dome positive population between *Bx^7^* mutants and control (Figure 2F & G), suggesting that progenitor pool is unaffected. However, staining with P1, a marker of differentiated plasmatocytes, revealed a marked reduction in differentiated plasmatocytes in *Bx^7^* lymph gland (Figure 2H & I).

Similarly, *Beadex* knockdown in medullary zone (MZ), cortical zone (CZ) and posterior signalling centre (PSC) leads to a comparable reduction in P1 positive differentiated plasmatocytes (Figure 3A-E). Surprisingly, in the intermediate zone we observed that *Beadex* knockdown leads to an increase in the plasmatocyte area, indicating that *Beadex* might prevent the premature differentiation of the progenitors in the intermediate zone of lymph gland (Figure 3F).

**Figure 3.**
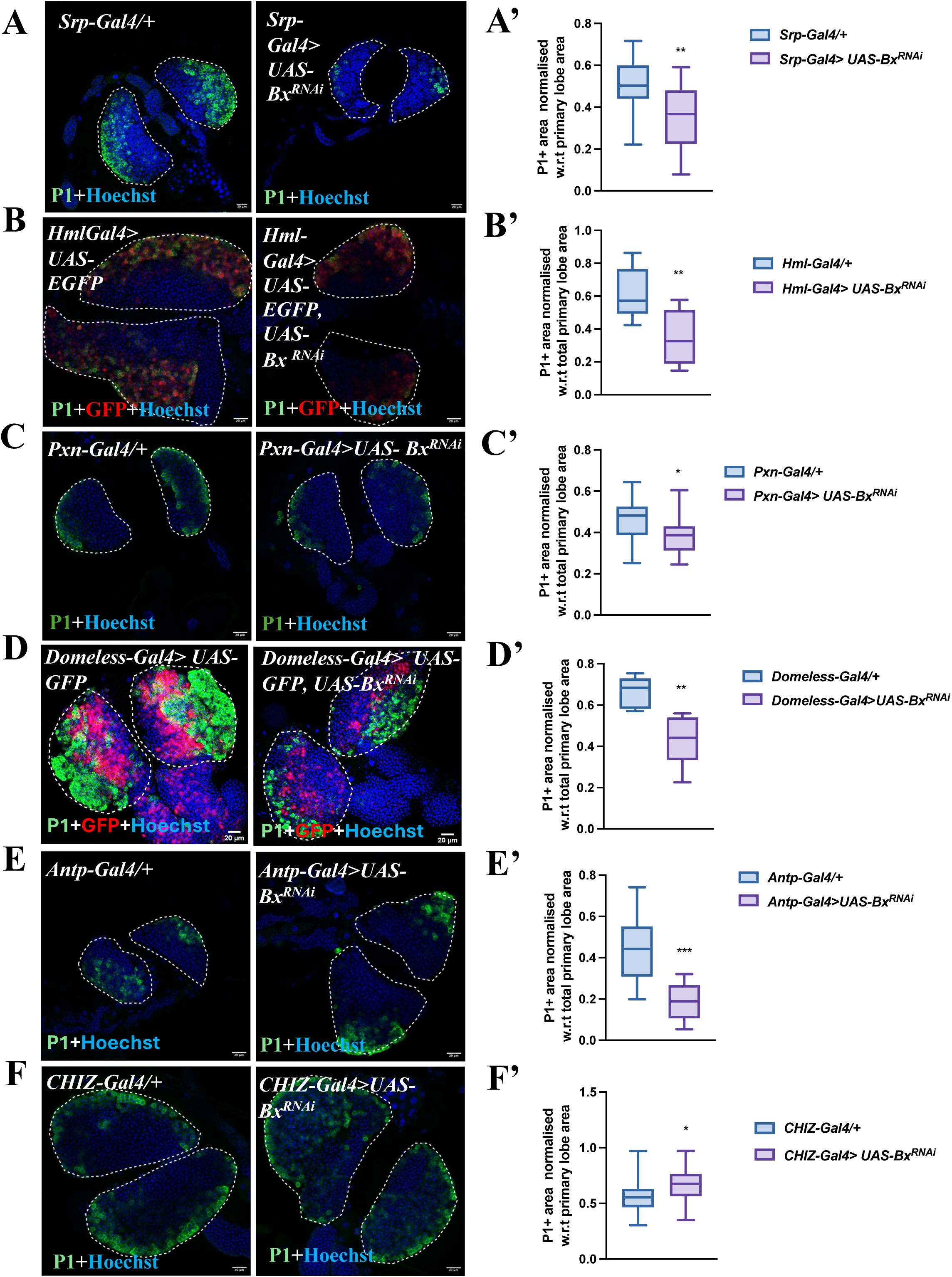
*Beadex* regulates plasmatocytes differentiation in zone specific manner. Confocal images of larval lymph gland primary lobe following Beadex knockdown in **A)** whole lymph gland using *Srp-Gal4*, **B)** in cortical zone (CZ) using *Hml-Gal4*, **C)** in mature plasmatocytes using *Pxn-Gal4*, **D)** in medullary zone (MZ) using *Domeless-Gal4*, **E)** posterior signalling centre using *Antp-Gal4* and, **F)** in the intermediate zone using *CHIZ-Gal4*. Differentiated plasmatocytes are marked by P1 (green) and nuclei using Hoechst (blue). The bar graphs on the right showed a significant reduction in the P1+ plasmatocyte area w.r.t total lymph gland area upon Beadex knockdown in whole lymph **A’)**, CZ **B’)**, mature plasmatocytes **C’)**, MZ **D’)** and PSC **E’)**. Beadex knockdown in IZ **F’)** showed a significant increase in P1+ area. The primary lobe of the lymph gland are demarcated by white dotted line. The graph represents the mean of three independent biological replicates and 20 lymph glands are analysed in each case. Statistical significance is denoted by asterisks: *p < 0.05, **p < 0.01.

### 3. *Beadex* role in phagocytosis is linked to actin cytoskeleton alterations

Plasmatocytes, the Drosophila equivalent of vertebrate macrophages, are the primary phagocytic cells responsible for eliminating pathogens and maintaining tissue homeostasis. Since *Beadex* mutants showed reduced plasmatocyte numbers, we next assessed for their phagocytic potential. In *Bx^7^* mutants, plasmatocytes exhibited a marked reduction in both the percentage of engulfing cells and the phagocytic index compared to controls (Figure 4A, A’, C, D). However, among the *Beadex* hypermorphic mutants, *Bx^J^*, the severe hypermorphic mutant (Figure 4H) displayed an increase in phagocytic index (Figure 4B, B’, B’’, J). A similar modulation of phagocytosis was observed upon hemocyte-specific knockdown of *Beadex* by RNAi (Figure 5A, B, D).

**Figure 4.**
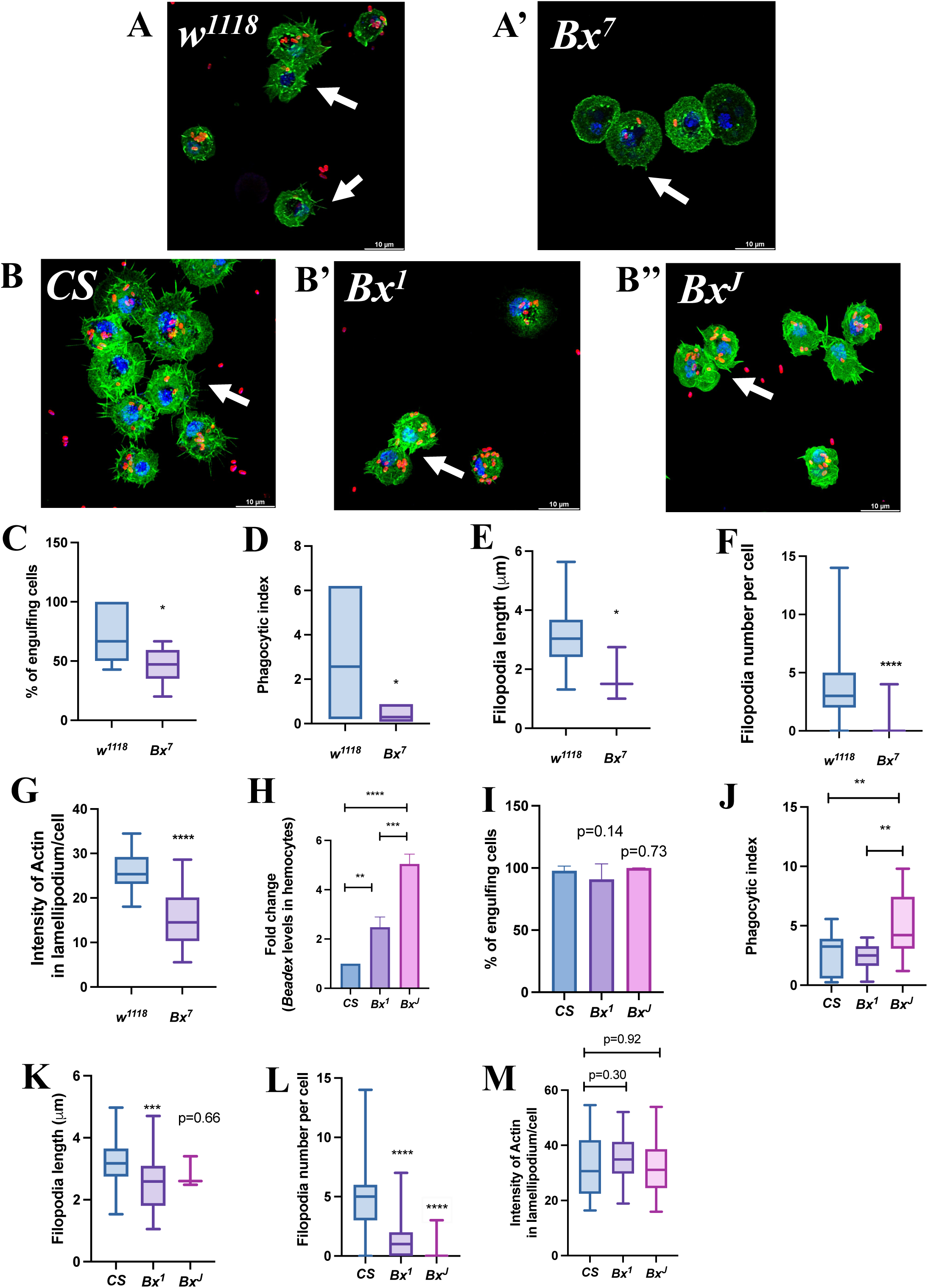
*Beadex* regulates phagocytosis and actin-based protrusions in dosage dependent manner. Confocal image of ex-vivo phagocytosis assay of plasmatocytes (green) isolated from **A)** control (*w^1118^*) and **A’)** *Bx^7^* mutant engulfing *E. coli* bioparticles (red). Cells were stained for F-actin (green) and nuclei (blue). Arrows indicate the filopodia extensions. Scale bar, 10 µm. Phagocytosis assay in hemocytes from **B)** CS controls, **B’)** & **B’’)** *Beadex* hypermorphic mutant (*Bx¹ & Bx^J^*). Phagocytosis and actin morphology are visualized as described in (A). Scale bar, 10 µm. **C)** Quantification of the percentage of hemocytes actively engulfing *E. coli* bioparticles in *w¹¹¹⁸* and *Bx⁷* mutants, showing a significant reduction. **D)** Quantification of the phagocytic index (number of particles per cell) in *w¹¹¹⁸* and *Bx⁷* mutants, showing a reduction. **E, F & G)** Quantification of filopodia number and length and intensity of actin in lamellipodium per cell in *w¹¹¹⁸* and *Bx⁷* mutants, showing a reduction. **I)** Quantification of filopodia length in *CS*, *Bx¹*, and *Bx^J^* genotypes. **H)** Relative transcript levels of *Beadex* in *CS* control and *Bx^1^* & *Bx^J^* mutant larvae, confirming the increased expression of *Beadex* in hemocytes. **I)** Quantification of the percentage of engulfing cells in *CS*, *Bx¹*, and *Bx^J^* genotypes. **J)** Quantification of the phagocytic index in *CS*, *Bx¹*, and *Bx^J^* genotypes, showing no change in *Bx¹* mutant and an increase in *Bx^J^* mutant. **G)** Quantification of filopodia number per cell in *CS* and *Bx^1^* & *Bx^J^* mutants, showing significant reduction. **K, L & M)** Quantification of filopodia number and length per cell and intensity of actin in *CS, Bx¹*, and *Bx^J^*genotypes. The filopodia number and length is reduced in these mutants with no change observed in lamellipodium area. The graph represents the mean of three independent biological replicates and 10-20 fields are analysed in each case. Statistical significance is denoted by asterisks: *p < 0.05, **p < 0.01, ***p < 0.001, ****p < 0.0001.

**Figure 5.**
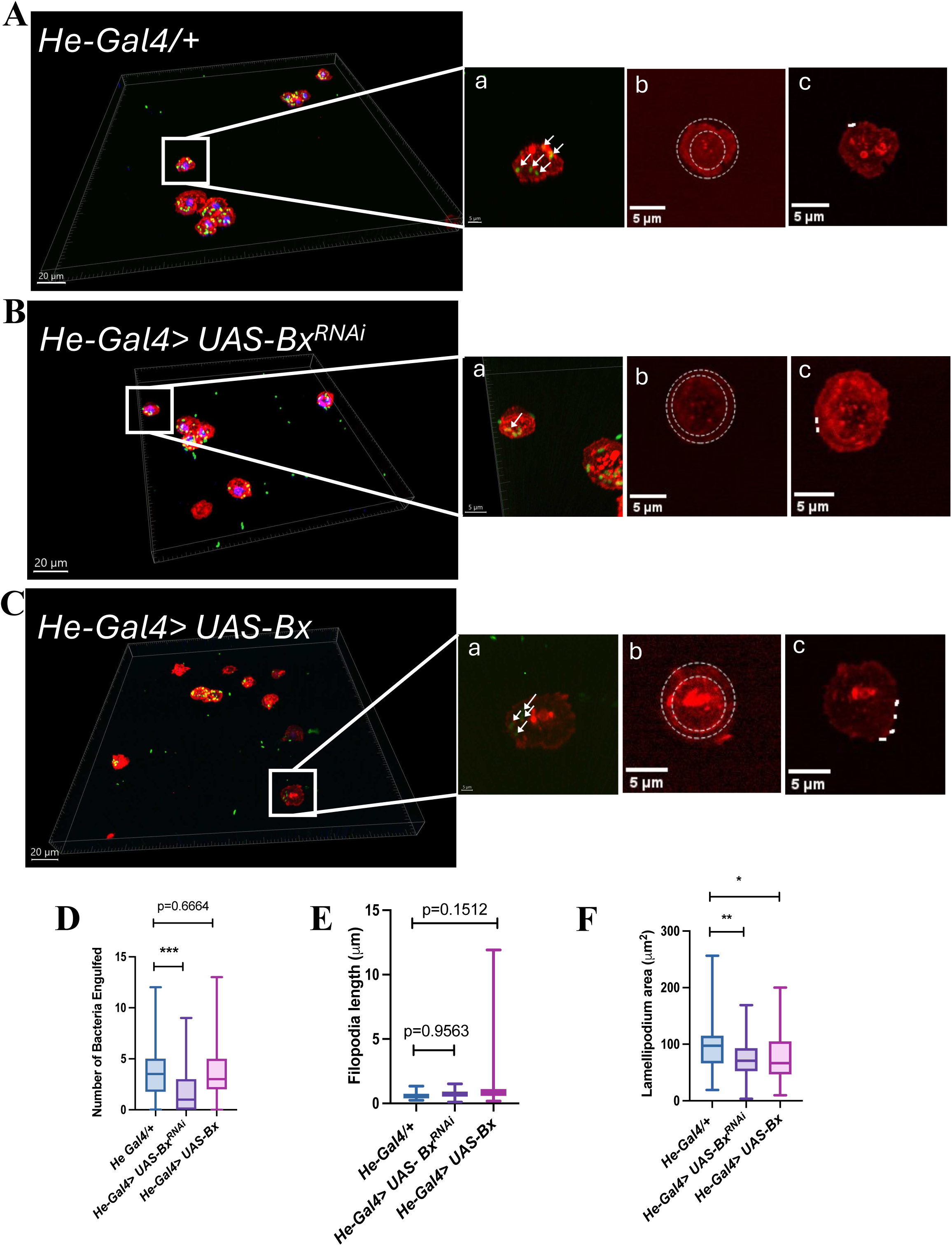
Both loss and gain of function of *Beadex* impairs phagocytosis and lamellipodium formation in hemocytes. Confocal microscopy images showing actin morphology and bacterial uptake in hemocytes with *He-Gal4* driven expression of **A)** control (*CS*), **B)** *Beadex* knockdown (*UAS*-*Bx^RNAi^*), and **C)** *Beadex* overexpression construct (*UAS-Bx*). Phalloidin staining is done to visualize actin filaments (red), and *E. coli* bacteria are shown in green. The leftmost images are 3D reconstructions. The smaller insets show magnified views of individual cells, demonstrating **a)** phagocytic cells, **b)** lamellipodia and **c)** filopodia in both knockdown and overexpression conditions. Quantifications of **D)** phagocytosis index, showing a reduction upon Beadex knockdown and no change upon Beadex overexpression in hemocytes, **E)** filopodia length & **F)** lamellipodium area in control, Beadex knockdown and overexpression hemocytes. Lamellipodia area is decreased with no observed change in filopodia length. The graph represents the mean of three independent biological replicates and 10-20 fields are analysed in each case. Statistical significance is denoted by asterisks: *p < 0.05, **p < 0.01, ***p < 0.001, ****p < 0.0001.

To understand the basis for these functional phenotypes of *Beadex* mutants, we examined the plasma membrane-associated cortical actin cytoskeleton, which is essential for forming the filopodia and lamellipodia required for particle engulfment. *Bx^7^* mutants (Figure 4E, F & G) plasmatocytes and the hemocyte specific *Beadex* knockdown plasmatocytes displayed a reduced number of filopodia and an overall decrease in lamellipodia area (Figure 5E & F).

Intriguingly, we observed a similar reduction in filopodia number and length in the hypermorphic mutants (*Bx^1^* and *Bx^J^*), despite their increased phagocytic efficiency (Figure 4K, L & M). These results indicate that Beadex modulates phagocytosis by modulating actin dynamics. However, in the hypermorphic mutants increased phagocytosis may not be driven by classical actin protrusions or caused because of some alternative mechanism such as altered membrane tension or a hyper-stable phagocytic cup, which could still manifest as a reduced lamellipodia area in our assays.

### 4. *Beadex* regulates the expression of Profilin, and Profilin overexpression rescues the phagocytosis defect

Phagocytosis begins with recognizing foreign molecules by receptors on the cell membrane, which activate various signalling pathways leading to activation of actin regulatory proteins. These proteins such as Ena, Fascin, Rho1, and Profilin, facilitates filopodia and lamellipodia formation. Given that Beadex is a known transcriptional co-regulator with nuclear localization, we hypothesized that it might directly affect the expression of these actin-associated genes.

To test this, we assessed the expression levels of actin regulators in *Beadex* loss-of-function and gain-of-function mutants. *Beadex* hypomorphic mutant (*Bx^7^*) showed a severe reduction in the levels of *profilin (chickadee*/*chic)*, which is an actin-monomer delivery protein with no changes observed in the bacterial recognition receptors (NimC1, SR-CI, Dscam1) (Figure 6A). Consistent with this, RNAi-mediated *Beadex* knockdown in hemocytes also resulted in a similar reduction in *profilin* expression (Figure 6E). Conversely, in the hypermorphic mutants (*Bx^1^* and *Bx^J^*), we observed a marked increase in the expression of *profilin* (Figure 6D), together with modest or no changes in NimC1, SR-CI, and Fascin (Figure 6B & C). However, in hemocyte specific overexpression of Beadex, a decrease in the profilin level was observed (Figure 6F).

**Figure 6.**
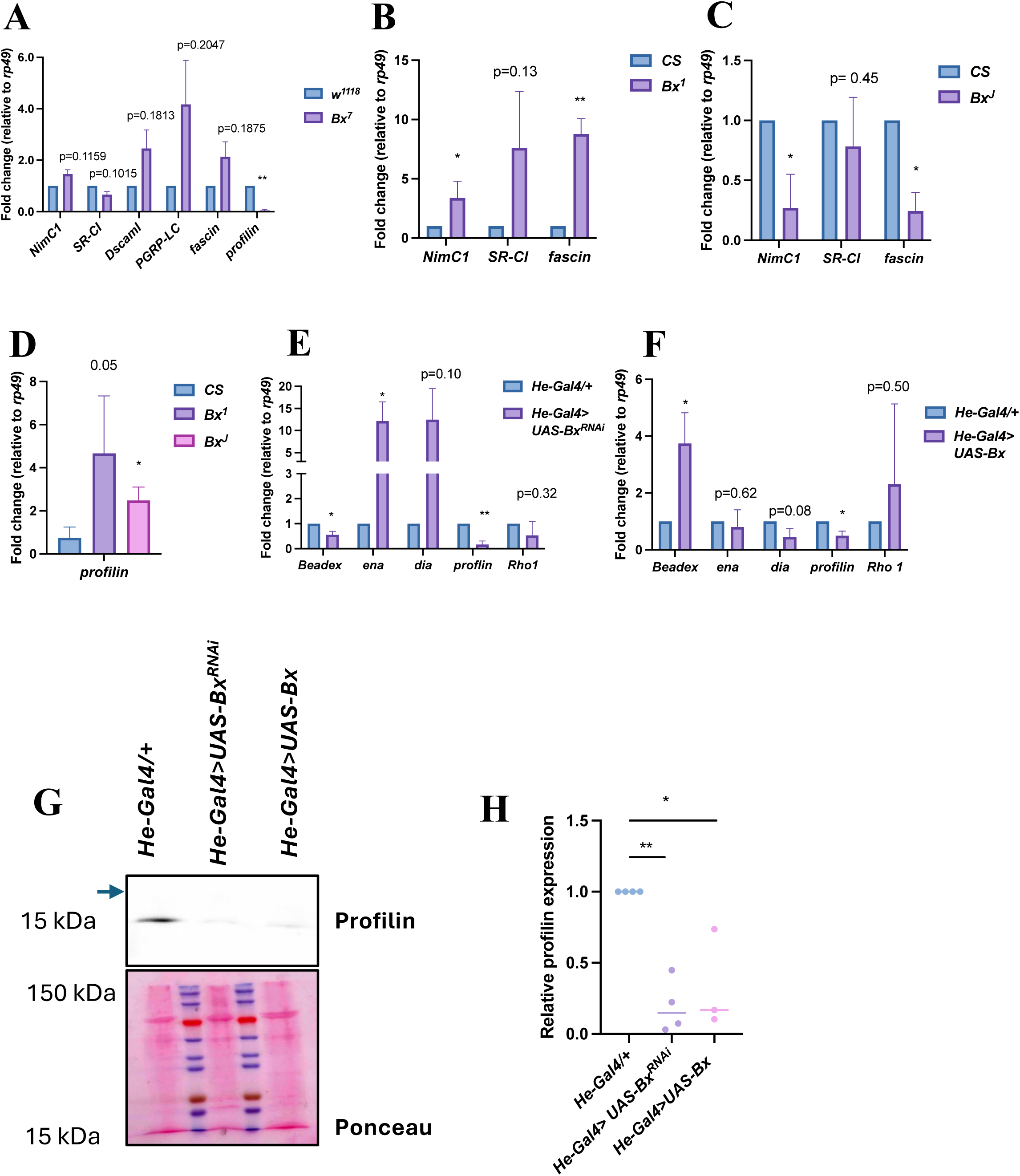
*Beadex* regulates actin regulated genes and protein expression. **A)**Relative gene expression level of receptors and downstream actin regulatory genes and hemocytes of *w^1118^* and *Bx^7^* mutants, showing a significant reduction in *profilin* expression level. **B & C)** Relative gene expression level of *NimC1*, *SR-CI* and *Fascin* in hemocytes of *Beadex* hypermorphic mutants (*Bx^1^* & *Bx^J^*). **D)** Relative gene expression of profilin in *CS*, *Bx^1^* and *Bx^J^* mutant showing a significant increase in hypermorphic mutants. **E & F)** Relative gene expression levels of several actin regulators in *He-Gal4>CS* controls and *Beadex* knockdown (*He-Gal4>UAS-Bx^RNAi^*) and overexpression (*He-Gal4>UAS-Bx*) in hemocytes, confirming the successful knockdown and overexpression of *Beadex* and the downregulation of *profilin*. **G)** Western blot analysis of Profilin (Chic) protein levels in hemocytes from *He-Gal4 > CS* controls, *He-Gal4 > UAS-Bx^RNAi^* knockdown, and *He-Gal4 > UAS-Bx* overexpression flies. **H)** Quantification of the Western blot in (E), showing a significant reduction in Profilin protein upon both Beadex knockdown and overexpression in hemocytes. The graph represents the mean of three independent biological replicates. Statistical significance is denoted by asterisks: *p < 0.05, **p < 0.01, ***p < 0.001, ****p < 0.0001.

This was corroborated at the protein level by immunoblotting wherein reduced Profilin levels upon *Beadex* knockdown and overexpression were observed (Figure 6G & H).

To confirm that reduced Profilin levels are the cause of phagocytosis defect, we have overexpressed *profilin (chic)* in the *Beadex* knockdown background (*He-Gal4> UAS Bx^RNAi^/UAS-Chic*) (Figure 7A, B & C). Overexpression of *profilin* in the *Beadex* knockdown background was sufficient to rescue the phagocytosis defect, confirming that reduced Profilin levels are the underlying cause of the phenotype (Figure 7E).

**Figure 7.**
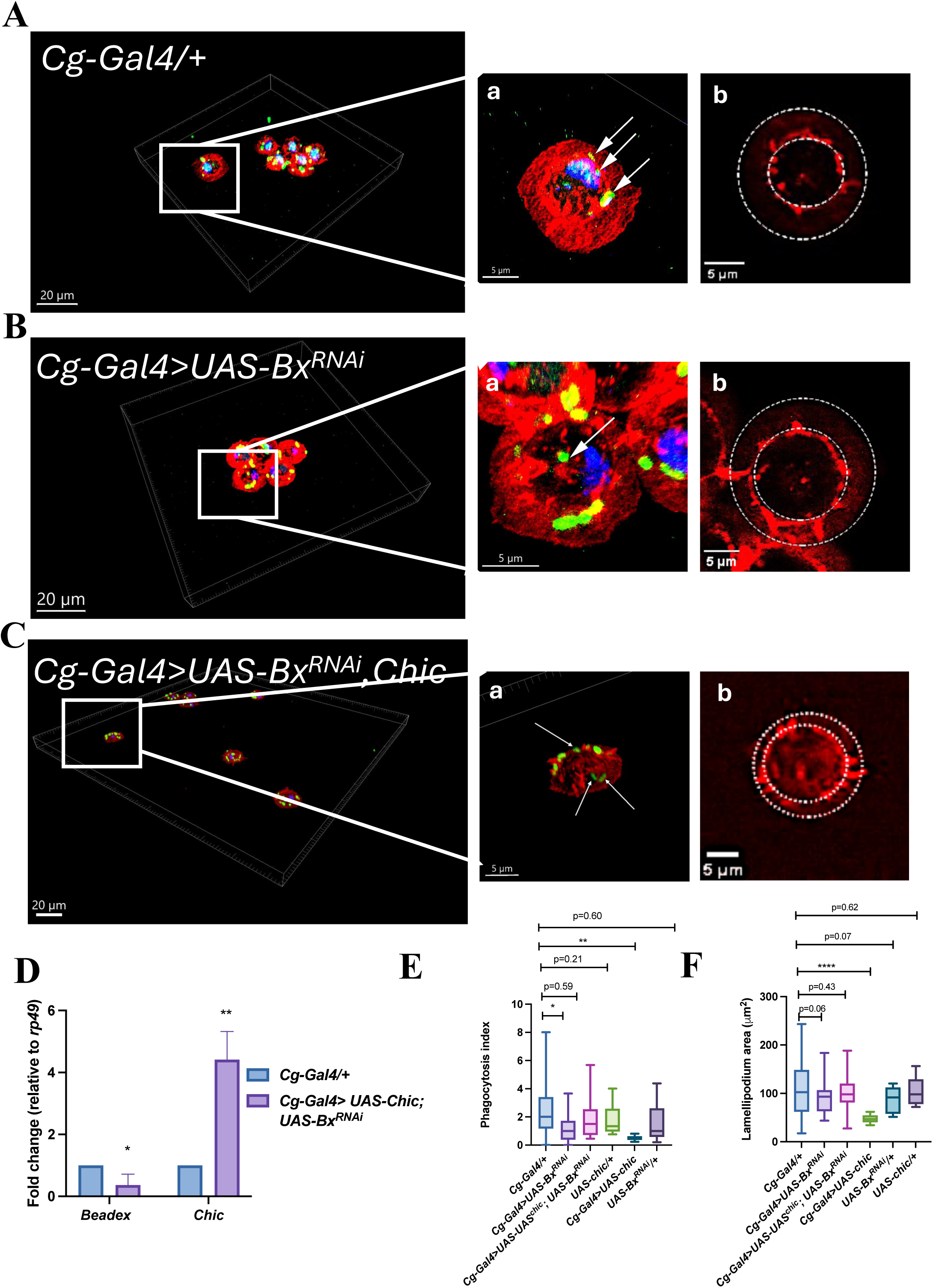
Overexpression of profilin rescues the phagocytosis defect of *Beadex* knockdown hemocytes. Confocal images of hemocytes showing actin morphology and bacterial uptake in respective genotypes: **A)** Control (*Cg-Gal4>CS*), **B)** *Beadex* knockdown (*Cg-Gal4>UAS-Bx^RNAi^*), **C)** profilin or chic rescue (*Cg-Gal4>UAS-Bx^RNAi^;UAS-Chic*). The cells are stained with phalloidin (red), *E. coli* (green) and nuclei is marked using Hoechst (blue). The insets **(a & b)** shows magnified images of individual cells. Engulfed bacteria are showing using white arrow and dashed lines are used to encircle lamellipodium. **D)** Relative gene expression levels of *Beadex* and *Chic* in control and *Chic* overexpressing hemocytes, confirming the successful overexpression of the *Chic* gene. **E)** Quantification of the phagocytosis index across the indicated genotypes. The data show a significant reduction in the phagocytic index in the *Beadex* knockdown condition, which is fully rescued by the overexpression of *Chic* (p=0.59). **E)** Quantification of the lamellipodium area across the indicated genotypes. The data show a significant reduction in lamellipodium area in the *Beadex* knockdown condition, which is also fully rescued by the overexpression of *Chic* (p=0.43). The graph represents the mean of three independent biological replicates and 10-20 fields are analysed in each case. Statistical significance is denoted by asterisks: **p < 0.01, ****p < 0.0001.

### 5. *Beadex* overexpression in hemocytes increases susceptibility to *Salmonella enterica serovar* Typhimurium infection independent of bacterial burden

Phagocytosis by immune cells plays a critical role in mounting an immune response against bacterial infections. To investigate whether Beadex influences immune defense, we assessed for the infection susceptibility of *Beadex* mutants following infection with the gram-negative intracellular pathogen *Salmonella enterica serovar* Typhimurium and *E. coli* as control (Figure 8A, B & G). We found that overexpression of *Beadex* in hemocytes using *He-Gal4* (*He*-Gal4*> UAS-Bx*) significantly increased fly susceptibility, leading to a dramatic reduction in survival post-infection upon *Salmonella* infection (Median survival of infected hemocyte specific *Beadex* overexpressing flies=7.5 (dpi), the p-value of Log Rank test: infected *He-Gal4/+* vs. *He-Gal4> UAS-Bx* is <0.0001) (Figure 8B). In contrast, hemocyte-specific knockdown of *Beadex* (*He-Gal4> UAS-Bx^RNAi^*) did not affect survival, suggesting that loss of *Beadex* function can be compensated for, or that its role in phagocytosis alone is not a primary determinant of survival against these pathogens.

**Figure 8.**
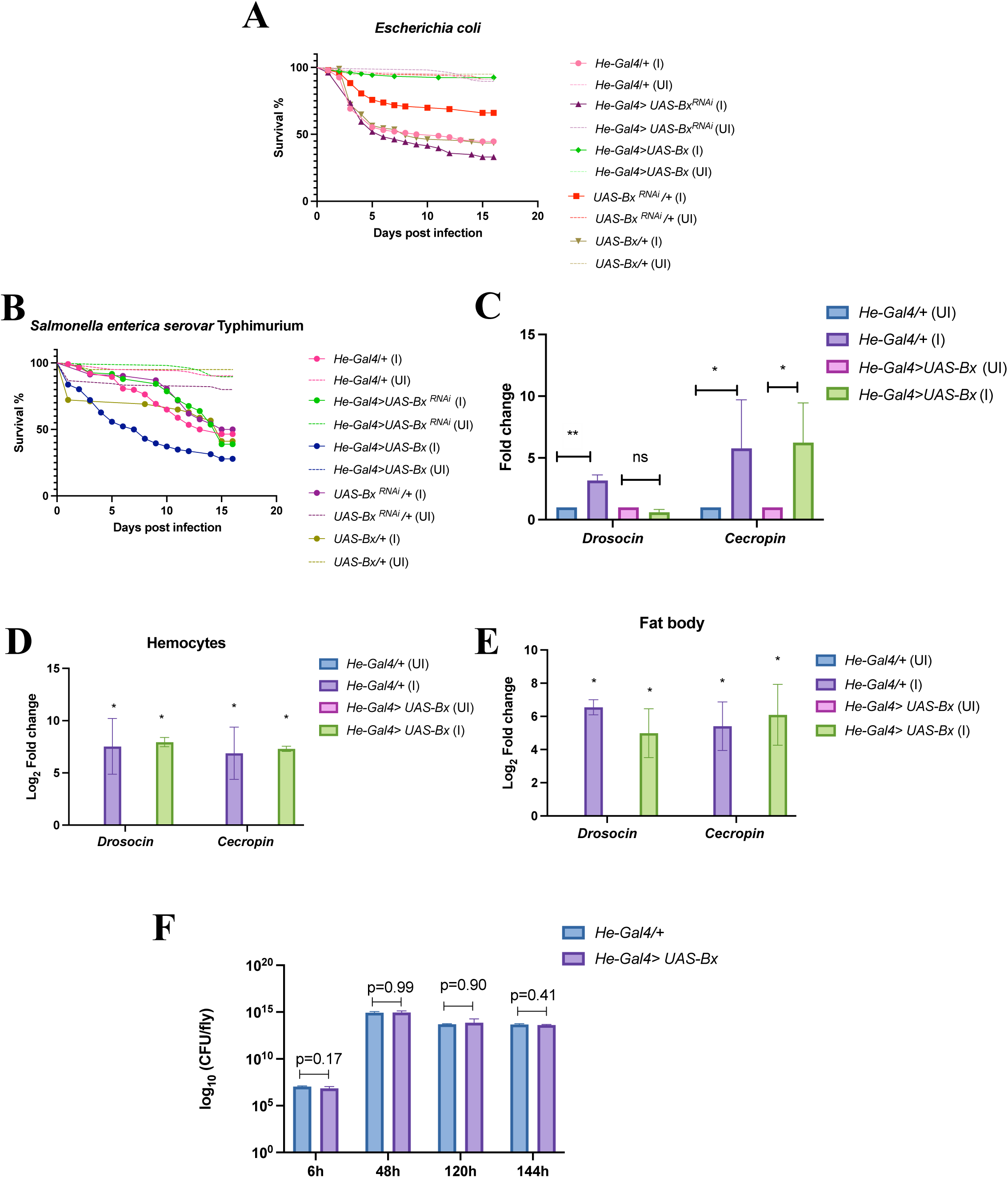
*Beadex* overexpression impairs host survival but not bacterial clearance. **A)** Survival curve of control flies (*He-Gal4/+)* and Beadex knockdown (*He-Gal4 >Bx^RNAi^*) and overexpression (*He-Gal4 >UAS-Bx*) genotypes following infection with *Escherichia coli*. and **B)** *Salmonella enterica* serovar Typhimurium. **C)** Relative gene expression levels of the antimicrobial peptides (AMPs) Drosocin and Cecropin in whole flies 6 hours after *Salmonella* infection. The data show significantly reduced levels of AMPs in the *Beadex* overexpression condition. **D)** Relative gene expression levels of Drosocin and Cecropin in hemocytes & **E)** fat body from control and *Beadex* overexpressing flies after *Salmonella* infection. **F)** Bacterial load (Colony Forming Units, CFU) in control and *Beadex* overexpressing flies measured at different time points after *Salmonella* infection, showing no significant difference between the two genotypes. For survival data of 100 flies are analysed in each genotype and bar graphs represents the mean of three independent biological replicates. Statistical significance is denoted by asterisks: *p < 0.05, **p < 0.01, ***p < 0.001. ns, not significant.

Given the severe susceptibility of *Beadex* overexpressing flies, our initial hypothesis was that these flies were unable to effectively control the bacterial infection. We therefore assessed the levels of antimicrobial peptides (AMPs) and measured the bacterial load by CFU after infection. To our surprise, while the AMP levels in the whole fly were not significantly changed in hemocytes specific *Beadex* overexpressing flies, we found that both hemocytes and the fat body produced AMPs at levels equivalent to controls (Figure 8C, D & E). Furthermore, the CFU counts in *Beadex* overexpressing flies after 6 days of infection were similar to those in control flies (Figure 8F).

These findings indicate that the heightened susceptibility of *Beadex*-overexpressing flies is not due to an inability to clear bacteria. Instead, overexpression likely triggers infection-associated pathology through mechanisms independent of bacterial burden, potentially involving dysregulated immune signalling or a harmful hyper-inflammatory response.

### 6. *Beadex* regulates a core transcriptional network regulating cytoskeleton components, immune and metabolic pathway in hemocytes

Our analysis has depicted that Beadex regulates phagocytosis by regulating the expression of key actin regulatory genes. This function is further supported by protein-protein interaction studies done in Drosophila hemocyte derived S2 cell line, which showed that Beadex interacts with transcriptional regulators and SWI/SNF chromatin remodelling complex proteins, BAP60 and Osa.

Therefore, to gain a comprehensive understanding of this complex regulatory network, we have performed a bulk RNA sequencing of larval plasmatocytes to investigate the molecular consequences of *Beadex* knockdown in hemocytes using the *He-Gal4* driver. This approach is thought to recapitulate the transcriptional changes of *Beadex* loss-of-function mutant (*Bx^7^*), thereby provide a comprehensive view of Beadex regulatory network in hemocytes.

Among the total 17895 genes analyzed, 1063 are differentially expressed with p-value <0.05 and log2FC of 1 between *Beadex* knockdown and control hemocyte samples. Within these differentially expressed genes, 595 genes are upregulated, and 468 are downregulated. Principal Component Analysis (PCA) revealed a clear separation between knockdown and control samples, suggesting a distinct transcriptional response. The most significantly upregulated genes include *PGRP-SC2, Impl2, Drsl2, Dgp-1,* and *LpR1*, whereas *CecropinB, CG31343,* and *CG31233* were among the downregulated genes (Figure 9A). Pathway-level analyses revealed that Beadex knockdown broadly affects immune signalling and metabolic regulation. GSEA highlighted significant downregulation of immune-related pathways, including “Immune System” (NES = –1.86, FDR = 0.047) and “Innate Immune System” (NES = – 1.56, FDR = 0.075), while enrichment analysis pinpointed specific processes such as “Phagosome” (KEGG dme04145) and “Antibacterial Humoral Response” (GO:0019731). These transcriptional changes directly corroborate our functional observations of impaired phagocytosis in Beadex mutants, linking Beadex to the regulation of vesicle trafficking and pathogen uptake machinery. Additionally, suppression of “Metabolism of Proteins” (NES = –1.68, FDR = 0.064) and enrichment of lipid and amino acid transport pathways indicate Beadex knockdown induces metabolic rewiring in hemocytes.

**Figure 9.**
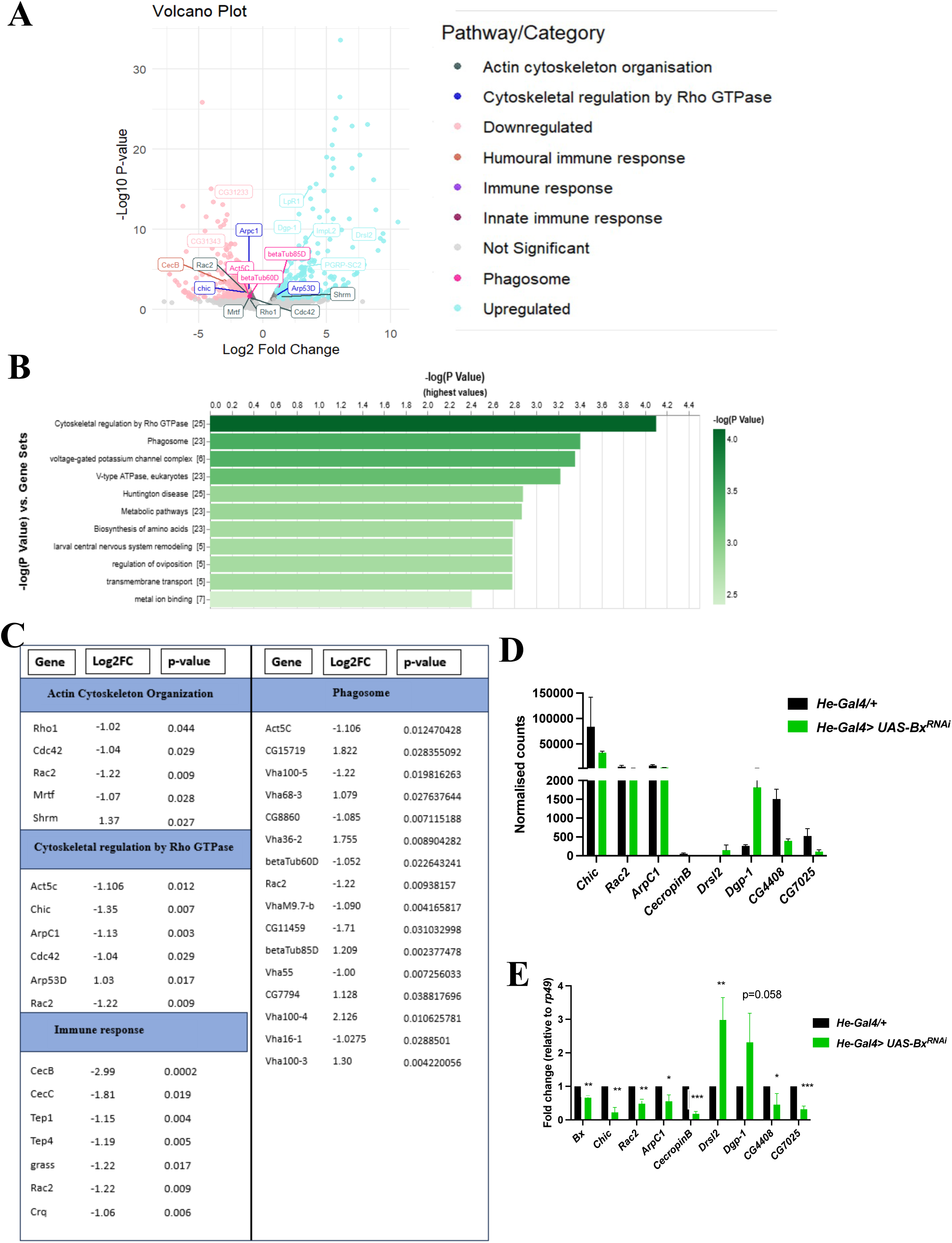
*Beadex* knockdown perturbs a transcriptional network regulating cytoskeleton and immune pathways. **A)** Volcano plot of differentially expressed genes following *Beadex* knockdown in larval hemocytes. Each dot represents a single gene, with colors highlighting specific enriched pathways. Genes with a log₂ Fold Change (log₂FC) > 1 and a p-value < 0.05 are considered differentially expressed. **B)** A bar chart showing the top significantly enriched pathways and gene sets, ranked by statistical significance (-log₁₀P value), including ‘Cytoskeletal regulation by Rho GTPase’, ‘Phagosome’, and ‘Immune response’. **C)** Tables listing a subset of differentially expressed genes within the most significantly enriched pathways, along with their corresponding log₂FC and p-values. **D)** Normalized RNA-seq read counts for a selected set of genes in control (*He Gal4 > CS*) and *Beadex* knockdown (*He Gal4 > BxRNAi*) hemocytes. **E)** Validation of RNA-seq data by quantitative PCR (qPCR) for the selected gene set, confirming the differential gene expression observed in the transcriptome analysis. Statistical significance in qPCR results is denoted by asterisks: *p < 0.05, **p < 0.01, ***p < 0.001.

Protein–protein interaction network analysis revealed strong enrichment of cytoskeletal components, including microtubule-based processes (GO:0007017; –log₁₀P = 15.5) and motor protein complexes (KEGG dme04814; –log₁₀P = 19.6). Enrichment of phagosome-associated (KEGG dme04145; –log₁₀P = 19.7) and antigen presentation pathways (R-DME-983170; –log₁₀P = 13.2) further supported a functional link between *Beadex* and phagocytic machinery (Figure 9B). Consistently, gene set enrichment analysis via PANGEA revealed significant enrichment of Cytoskeletal regulation by Rho GTPase, V-type ATPase, Voltage-gated potassium channel complexes, and metabolic modules including amino acid biosynthesis and the citrate cycle (Figure 9C). qRT-PCR validation of selected differentially expressed genes showed the similar trend as observed in the transcriptomic analysis (Figure 9D & E).

Together, these converging analyses implicate *Beadex* as a key regulator of immune function and metabolism in Drosophila hemocytes, potentially acting through cytoskeletal remodelling, intracellular trafficking, and phagocytic control. The transcriptional changes we observed are consistent with Beadex acting as broad regulator of gene expression, linking in concert with SWI/SNF chromatin remodelling complex. Despite these changes, the absence of overt infection susceptibility in knockdown flies suggests the existence of compensatory immune mechanisms, warranting further investigation.

## Discussion

Innate immune cells are fundamental in maintaining tissue homeostasis by either regulating the cellular immune response required to fight pathogen or regulating tissue integrity by clearing the necrotic and apoptotic debris. The cellular process central in this immune cell surveillance is phagocytosis and actin cytoskeleton plays an essential role in regulating it(Mylvaganam et al., 2021). In this study, we uncover a novel role of Drosophila LIM-only protein, Beadex, in regulating the phagocytosis by regulating the expression of several actin cytoskeleton regulatory proteins. Notably, we find that *Beadex* controls the expression of *profilin* (encoded by *chickadee*), a master regulator of actin filament turnover that also influences proliferation, apoptosis, and oncogenic signalling (Guruharsha et al., 2012). Profilin helps in actin polymerisation by replenishing the pool of activated actin monomers by facilitating the exchange of ADP for ATP on actin monomers. Previous studies have shown that changes in the levels of profilin impact overall actin levels in the cells. Increased profilin can also change the balance between free G-actin and polymerized actin and can have a complex effect on the kinetics of filopodia and lamellipodia formation (Pernier et al., 2016). Therefore, the decrease in the phagocytosis rate could be because of this imbalance between actin polymerization and depolymerization rates due to changes in the profilin levels in the cells.

Importantly, Profilin dysregulation impacts hematopoiesis, as reported in *chickadee* overexpression mutants, which exhibit an abnormal increase in plasmatocyte numbers, reduced lamellipodium area, and compromised phagocytic potential. Given Profilin’s nuclear functions, where it interacts with transcriptional machinery, Beadex may regulate cytoskeletal remodelling through both direct transcriptional control and secondary effects of Profilin on gene expression, reinforcing its role as a transcriptional hub for actin remodelling.

Interestingly, while LMO2, the mammalian ortholog of Beadex has been reported to promote breast cancer cells metastasis through cytoplasmic interactions with Arp2/3 complex and Profilin (Liu et al., 2017), we find that Beadex in Drosophila hemocytes predominantly localizes to the nucleus, pointing toward a transcriptional regulatory role. Nuclear localization of LMO2 was also observed in mouse bone marrow–derived macrophages and human THP-1 cells (unpublished data, UN Lab), suggesting a conserved immune-specific nuclear function of LMO2-family proteins distinct from their motility-promoting roles in epithelial cancers. This context-dependent subcellular localization may underlie their functional versatility across cell types.

RNA sequencing of *Beadex* knockdown hemocytes, reveals a number of genes that are perturbed upon *Beadex* knockdown in hemocytes which includes downregulation of anti-microbial peptides (*Cecropin*), phagocytic receptor (*Croquemort*), regulators of vesicle trafficking and several RhoGTPases including *Cdc42*. These changes reflect the phagocytosis defects observed in vivo and underscore *Beadex’s* central role in immune defense. Protein–protein interaction studies in S2 cells further support this model, identifying Beadex associations with BAP60 and Osa, components of the SWI/SNF chromatin remodelling complex (Guruharsha et al., 2012). Given the SWI/SNF complex’s established role in coordinating immune and metabolic transcriptional programs, Beadex likely functions as a LIM-domain scaffold that recruits chromatin remodelers to cytoskeletal and immune effector gene loci, accounting for the >1,000 differentially expressed genes observed in our transcriptomic analysis.

In addition, our findings suggest that *Beadex* overexpression in hemocytes may disrupt immune homeostasis. Intracellular pathogens like *Salmonella enterica serovar* Typhimurium are largely restricted to hemocytes, where host trigger additional immune responses to contain infection upon release of bacterial effector molecules(Brandt et al., 2004). In flies overexpressing *Beadex*, mortality occurs despite bacterial loads comparable to controls, suggesting that death may result from excessive cytokine signalling rather than uncontrolled bacterial proliferation. In Drosophila, hemocytes can secrete cytokine-like molecules, such as the TNF homolog Eiger and Unpaired (Upd) ligands, to amplify systemic immune responses. We propose that *Beadex* overexpression may enhance cytokine release from hemocytes, effectively restricting bacterial growth but causing collateral tissue damage and systemic stress, ultimately leading to host death.

Functional parallels with vertebrate hematopoiesis further strengthen this model. In vertebrates, LMO2 forms a complex with LIM-binding protein, Ldb1, and a recent study has shown that this complex regulates the maintenance of hematopoietic stem cells (Cleveland et al., 2013). In acute myeloid cell lines, it is shown that LMO2 and Ldb1 are mutually regulated. LDB1 regulates the proliferation and apoptosis of AML cell lines and the expression of multiple genes, including LMO2. In the absence of LMO2, the proliferation of AML cell lines is drastically reduced, and the cells undergo apoptosis(Lu et al., 2023a). The S2 cell interactome data has also shown the interaction of *Beadex* with *Chip* (fly homolog of LDB1) in Drosophila, hinting towards a similar mechanism of action(Guruharsha et al., 2012; Milán et al., 1998b).

Our findings also extend prior work from our lab showing *Beadex’s* role in crystal cell specification via modulation of Pannier (Pnr), a GATA transcription factor central to lymph gland development. STAT signalling plays an important role in regulating *pannier* (*pnr*) transcription levels in the cortical zone in the lymph gland to regulate plasmatocyte development(Minakhina et al., 2011). Unidentified additional factors regulate *pannier* expression in the third instar larval lymph gland. Moreover, ectopic overexpression of *pannier* in hemocytes has an inhibitory effect on lymph gland growth, particularly CZ expansion and plasmatocyte development(Minakhina et al., 2011). As *Beadex* is a known genetic interactor of *pannier*, it is plausible that via *pannier*, *Beadex* regulates the lymph gland size and plasmatocyte development.

Together, these findings establish Beadex as a key nuclear regulator linking chromatin remodelling and cytoskeletal gene transcription to immune cell function. By revealing a conserved, context-specific role of LMO2-family protein, Beadex in innate immunity in Drosophila, our study highlights how transcriptional scaffolds can dynamically integrate cytoskeletal remodelling, differentiation, and effector responses. This mechanistic framework provides new insights into LIM-only protein biology and offers potential targets for modulating phagocyte function in health and disease.

## Materials and Methods

### Drosophila melanogaster strains and maintenance

Flies were reared on cornmeal agar medium, maintained on a 12-hour day/night cycle at 25°C unless specified otherwise. All knockdown and overexpression crosses were performed at 29°C. Following fly lines were used: *Canton-S* (BS#1), *Bx^7^* (Kairamkonda and Nongthomba, 2014), *Bx^1^* (Bloomington stock BS#15), *Bx^J^* (BS#3997), *He*-*Gal4* (BS# 8900), *UAS-Bx* (gift from Prof. S. M. Cohen, SMC, Singapore), *UAS-Bx^RNAi^* (VDRC# 2917), *Cg*-*Gal4* (BS#7011), *UAS-Chic* (kind gift from Prof. Stephane Noselli, iGV, France), Domeless-GFP (kind gift from Dr. Rohan J. Khadilkar, ACTREC, Pune), *Hml-Gal4* (BS#30140), *Pxn*-*Gal4* (BS#600223), *Bx*-*Gal4* (BS#84280), *UAS-Bx-RB FLAG-HA* (generated in Fly facility, NCBS, Bengaluru), *UAS-GFP*, *Antp*-*Gal4*, *Srp*-Gal4 and *CHIZ*-*Gal4* (kind gift from Prof. Maneesha Inamdar, JNCSAR, Bengaluru)

### Plasmatocyte counting

*Hml*-GFP was used as a reporter to count larval plasmatocytes. Each larva was placed in a drop of Schneider’s media (SM) on a glass slide, and the cuticle was pinched to release the hemocytes. The dissected larvae were transferred to the second drop of SM and scraped to remove the sessile hemocytes(Petraki et al., 2015).

In adult flies, *Hml*-GFP was used as a fluorescent reporter to mark plasmatocytes, and the number of GFP-positive cells in the thorax, three legs, and head of one side of the fly was counted as described previously(Boschet al., 2019).

### Ex-vivo phagocytosis assay

Hemolymph was collected from 5 third instar larvae of each strain*—He-*Gal4*/+, He-*Gal4*> UAS-Bx^RNAi^*, *He-*Gal4*> UAS-Bx*, *w^1118^,* and *Bx*^7^, by making a slit on the posterior side of the larvae in Schneider’s media containing 1 nM phenylthiourea using sharp needles. The collected hemolymph was then allowed to attach to the coverslips for 20 minutes, followed by a 1X PBS wash. Heat-killed *E. coli* (K-12) GFP-tagged bacteria pellet was resuspended in Schneider’s media, and 100 µl of it was added onto the coverslip kept on ice for 25 minutes for *He-*Gal4*/+, He-*Gal4*> UAS-Bx^RNAi^*, *He-Gal4> UAS ^-^Bx* and for *w^1118^*and *Bx*^7^ followed by 1X PBS wash. Warm Schneider’s media containing phenylthiourea (PTU) were added onto the coverslip, which was then kept at 29°C for 20 minutes, followed by a 1X PBS wash. The bacterial cell mixture was fixed by adding 4% PFA onto the coverslip for 10 minutes at room temperature, followed by a 1X PBS wash. Cells were then preincubated in PBST for 20 minutes and then incubated with Phalloidin (1:250, Phalloidin Atto 565, 94072, sigma) and Hoechst (1:1000) for an hour. After two 1X PBS washes, the coverslip was mounted using a drop of mounting media onto a slide, and the edges of the coverslip were then sealed. Images were obtained using Leica SP8 confocal system.

### Dissection and staining of lymph glands

Wandering third-instar larvae were staged properly by allowing the mated females to lay eggs on an egg laying plate for 12 hours. Following hatching the synchronised larvae were collected after every 3 hours. The lymph gland was dissected by placing the larvae on a drop of PBS on slide. Following dissection, the whole head complex containing brain and lymph gland attached to the mouth hook was taken and fixed in 4% PFA for 25 minutes, followed by 3 washes with PBSTx (1X PBS + 0.3% Triton X-100) for 15 minutes each. This is followed by blocking in 5% BSA followed by incubation with primary antibodies, anti-P1 (1.50, kind gift from Prof. Istvan Ando, HUN-REN, Hungary) and anti-Antp (1.100, 4C3, DHSB). Following overnight incubation, tissues were washed thrice and treated with secondary antibody for 2 hours at room temperature. Following incubation, tissues were again washed and mounted in the mounting media (propyl gallate and glycerol) and imaged using Leica falcon confocal microscope.

### Image analysis and quantification

Analysis was performed using ImageJ-win64 and ImarisViewer (10.2.0). Cell area was calculated by creating a Z-stack projection (max intensity), selecting the boundaries of the cell using the freehand selection tool, and then measured the area in ImageJ. The phagocytosis index was calculated by manually counting the ingested bacteria and viewing the cell in 3D in Imaris. The lamellipodia area was calculated by creating a Z-stack projection (max intensity), and then selecting the area with lamellipodia and without the lamellipodia using the freehand selection tool; the two values were then subtracted to get the lamellipodia area. Filopodia length and number were calculated by using the ImageJ plugin, semi-automated FiloQuant (Jacquemet et al., 2017). Phalloidin cumulative total cell fluorescence was calculated by creating a Z-stack projection (max intensity), then selecting the boundaries of the cell using the freehand selection tool and measuring the area and integrated density. The mean fluorescence of the background was calculated by using a rectangular selection tool to select a section of the background of the image that is devoid of any fluorescence signal, and then by measuring the mean grey value, an aggregate of five such different sections of the same image was used to calculate. Cumulative total cell fluorescence (CTCF) was calculated using the formula CTCF = Integrated Density - (Area of selected cell x Mean fluorescence of background readings).

For calculating the area of P1 positive cells in lymph gland, middle two stacks were taken, P1 stained area is marked using freehand tool in ImageJ and normalised with the total primary lobe area.

### Septic injury for infection

*Escherichia coli* (*E. coli*) and *Salmonella enterica serovar* Typhimurium were used for septic Gram-negative bacterial infections, and *Staphylococcus aureus* was used for septic Gram-positive bacterial infection. Bacteria were cultured overnight in LB-Ampicillin media, pelleted, and resuspended in 50 μL of sterile water. Flies were infected in the thorax using a sharpened tungsten needle dipped in the concentrated bacterial suspension or sterile water for control—survival rates of flies and colony-forming units (CFU) after the treatment were measured under identical conditions for all test genotypes. One prick with the tungsten needle is approximately 200 CFU/fly for *Salmonella enterica serovar* Typhimurium and, 292 CFU/fly for *E. coli*.

### Quantitative or real time PCR (qRT-PCR)

RNA was isolated using Trizol method of RNA isolation either from whole adult fly, adult fly fat body or hemocytes and from third instar larvae hemocytes. Briefly, for isolating fat body from adult flies, flies were placed in a drop of PBS, the abdomens were removed from the upper part of the body, opened to remove the gut, ovaries and other organs and cuticle attached with the fat tissue were collected and stored in Trizol reagent.

For collecting hemocytes from adult flies, 20 flies were taken in a 0.6 ml Eppendorf tube in which a hole is created at the bottom to collect the haemolymph and placed in a 1.5 ml Eppendorf tube. These flies were then covered in glass beads and centrifuged at 10000 rpm for 20 minutes at 4°C. This was repeated twice for a total of 40 flies and the collected hemolymph was dissolved in 300 µl of Trizol reagent.

For collecting hemocytes from larvae, they were placed in a drop Schneider’s medium containing PTU and their cuticle was pinched using a needle to release the hemolymph. The collected hemolymph was centrifuged at 1000 g for 5 minutes at 4°C to collect the hemocytes, which was further dissolved in 300 *u*l of Trizol reagent.

RNA was isolated using manufacturer’s instruction and the quantity and quality of the isolated RNA were assessed using Nanodrop ND-1000 spectrophotometer. RNA was treated with 1U of DNase I (EN0521, Thermo scientific) to remove DNA contamination and converted into cDNA using Revertaid reverse transcriptase (6110A-PrimeScriptTM 1^st^ strand cDNA Synthesis Kit). qRT-PCR was performed using NEB Sybr mix on an applied biosystems QuantStudio™ 3 Real-Time PCR System. Expression values were normalised to control *rp49*.

### Analysis of whole genome mRNA expression in hemocytes by RNA sequencing

RNA sequencing analysis was performed on 2 biological replicates of the test (*He-Gal4*> *UAS-Bx^RNAi^*) and control (*He-Gal4*> *CS*) samples. The RNA was isolated from approximately 200 larval hemocytes as described above, using Trizol method of RNA isolation. The isolated total RNA was quantified using Nanodrop. The integrity of RNA was evaluated on 1% agarose gel. Raw reads were checked for base quality and adapter content using FastQC (v0.12.1). Fastp (0.23.2) was used to remove adapter content and trim low-quality bases. The reads were mapped to the *Drosophila melanogaster* BDGP6.46 reference genome using STAR aligner (Dobin et al., 2013). The aligned genes were quantified using HTSeq (Anders et al., 2015). The genes showing differential expression between the RNAi samples and control samples were analyzed using the R package DESeq (Love et al., 2014). A volcano plot was generated using the R package EnhancedVolcano. The gene set enrichment analysis (GSEA) of differentially expressed genes with p-value <0.05 and log2FC of 1 between *Beadex* knockdown and control hemocyte samples was performed using the PANGEA (Pathway, Network and Gene-set enrichment analysis) tool (Hu et al., 2023). The RNA sequencing data have been deposited in the NCBI Sequence Read Archive (SRA) under BioProject accession number [PRJNA1338986].

### Western Blotting

Hemocytes were isolated from third-instar Drosophila larvae as described previously. The isolated hemocytes were resuspended in 1X RIPA buffer containing protease inhibitor cocktail and PMSF. The cells were lysed mechanically, and cell debris were cleared by centrifugation at 12000 rpm for 15 minutes. The protein concentration was estimated using Bradford protein assay (786-012 CB™ Protein Assay). For western blot analysis 50 µg of protein was denatured in 5X Laemmli buffer at 95°C for 5 minutes and separated on a 12% SDS-polyacrylamide gel. Following electrophoresis, the proteins were transferred to a PVDF membrane at 200 mA constant current for 2 hours. The membrane was stained with ponceau and then blocked in 5% BSA for 1 hour. The membrane was then incubated overnight at 4°C in primary antibody, anti-Chic (Chi 1J-DHSB-1.100). After washing with TBST, the membrane was incubated with the appropriate HRP-conjugated secondary antibodies for 1 hour at room temperature. Protein bands were visualized using an enhanced chemiluminescence ECL reagent (1705061, Bio-Rad) and detected with the ChemiDoc imaging system (Bio-Rad). The mean Gray value or intensity of bands were quantified in ImageJ and normalised with the total protein intensity observed following ponceau staining.

### Statistical analysis

All Statistical analyses were performed using GraphPad Prism 10.0. Student’s t-test was used to compare the mean of two groups, and One-way ANOVA and an appropriate post-hoc test (Dunnett’s multiple comparisons test) were used for more than two groups.

**Table 1.**
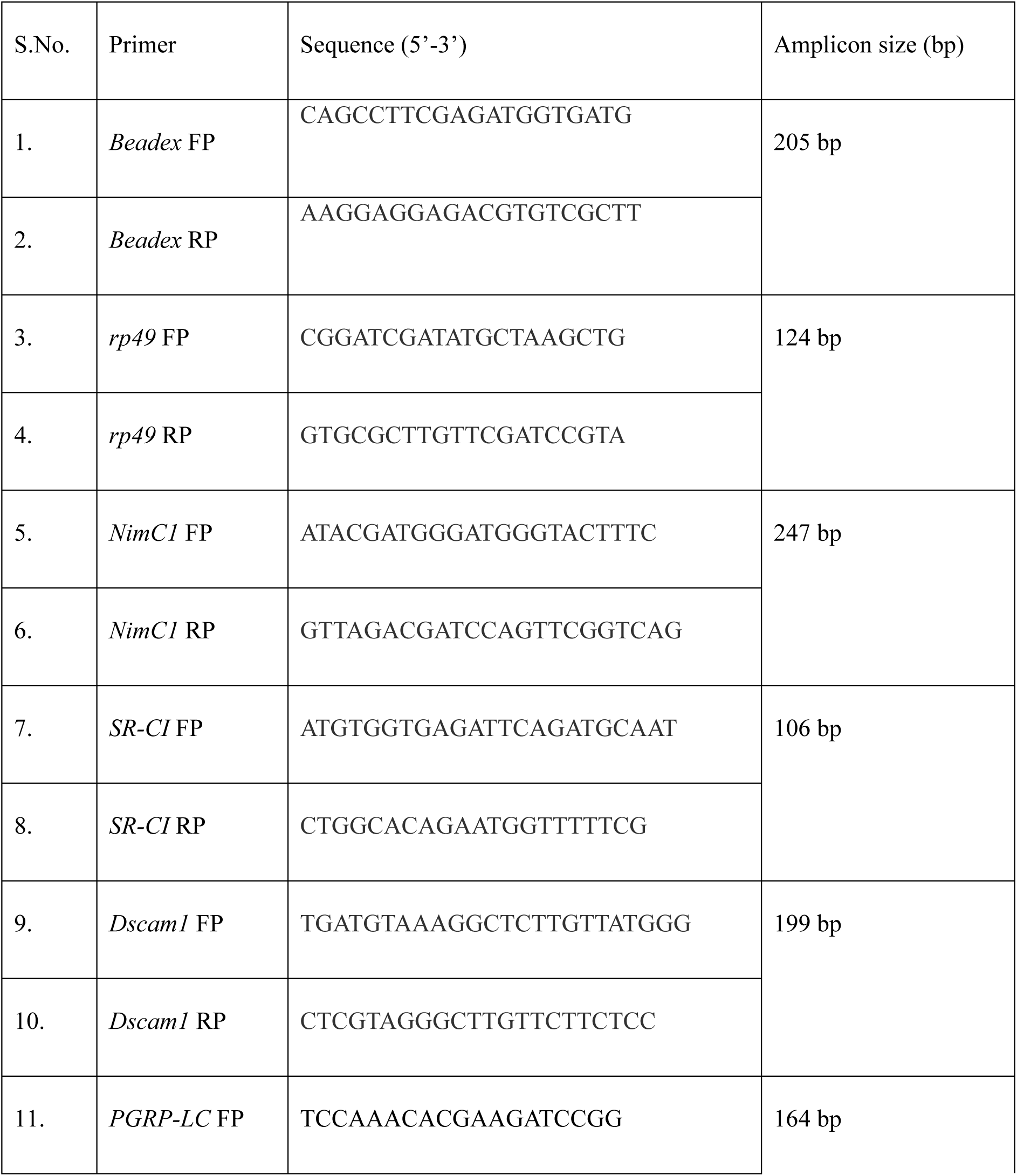
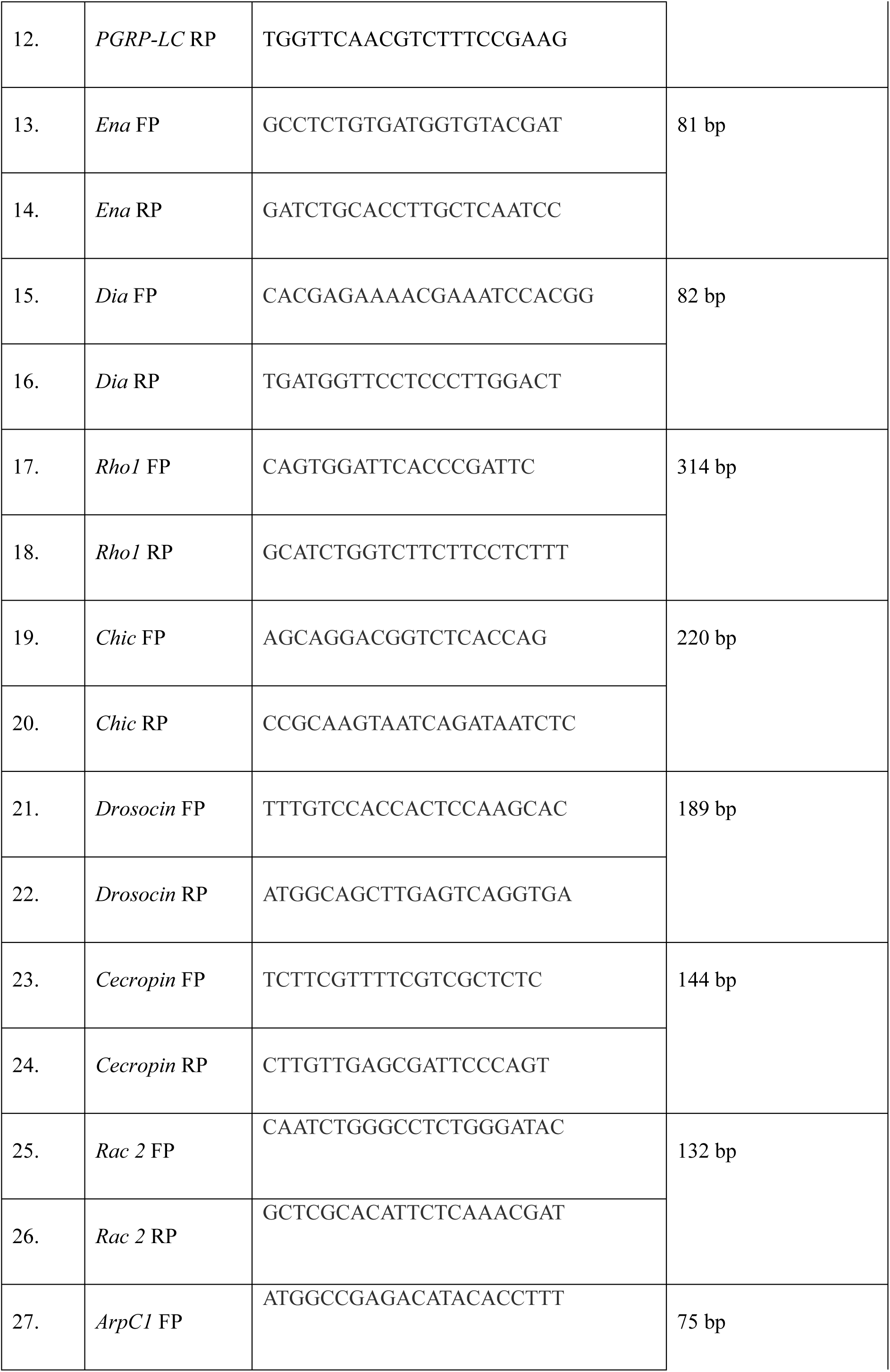
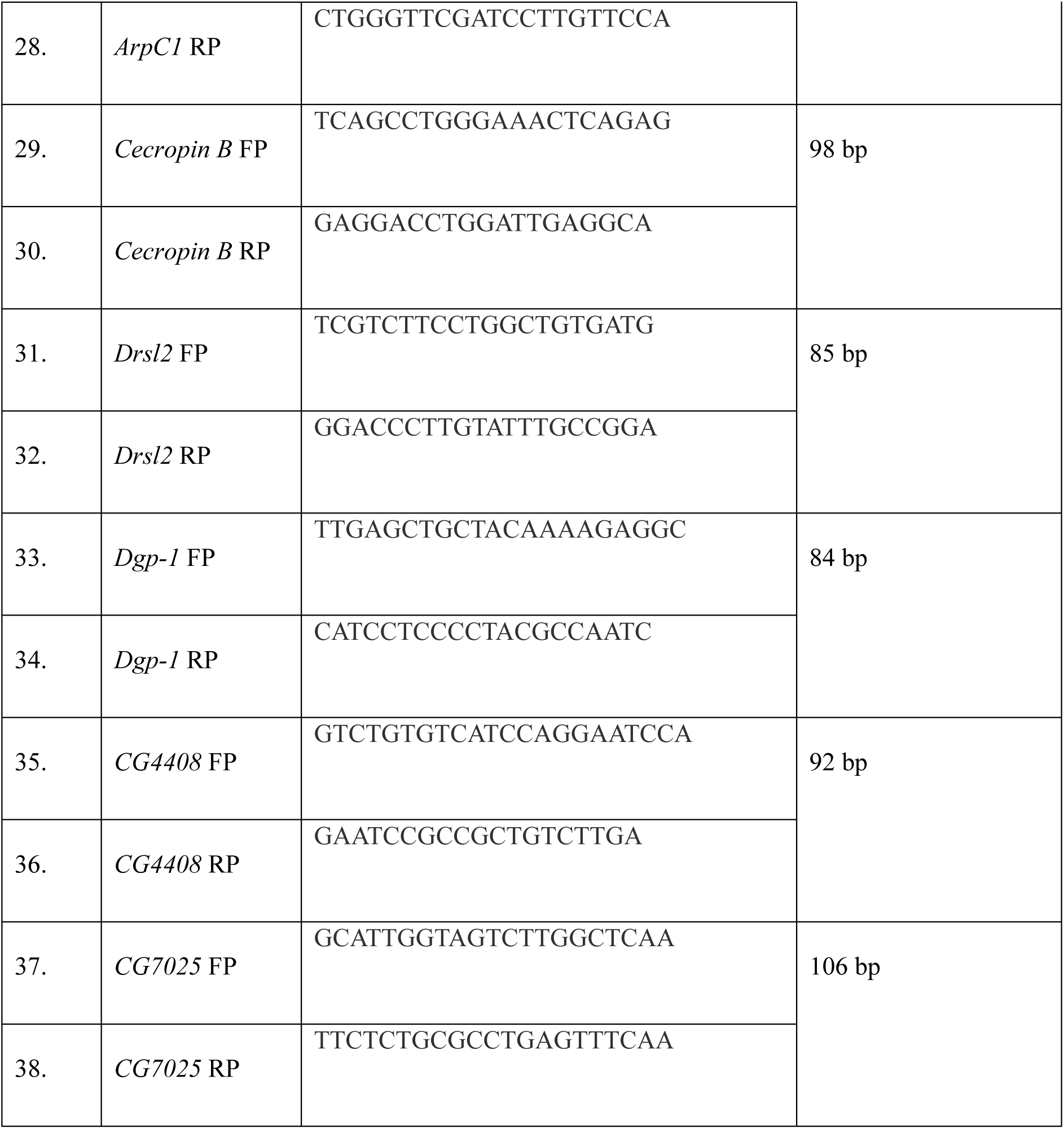
List of primers used in Quantitative or real time PCR (qRT-PCR):

## Author Contribution

Conceptualization, Funding acquisition, Project administration, Supervision, Resources: UN, SJ Methodology, Data curation, Formal Analysis, Validation, Visualization: SJ, KS, VS Manuscript Writing – review & editing: SJ, UN

## Consent for publication

All the authors have provided consent for publication.

## Data Availability

All primary data generated in this study are provided in the manuscript. Further inquiries or requests for additional information may be addressed to the corresponding author.

## Acknowledgments

We thank the Indian Institute of Science (IISc), Bangalore, for institutional support and research facilities. S.J. acknowledges the Prime Minister’s Research Fellowship (PMRF), Government of India, for financial support. We are grateful to Prof. S. M. Cohen (SMC, Singapore), Prof. Stéphane Noselli (iBV, France), Dr. Rohan J. Khadilkar (ACTREC, Pune), Dr. Tina Mukherjee (inStem, Bengaluru) and, Prof. Maneesha Inamdar (JNCASR, Bengaluru) for generously providing fly lines used in this study. We also thank the NCBS Fly Facility for generating transgenic lines and all members of our laboratory for valuable discussions and technical assistance.

